# Deuterated Polyunsaturated Fatty Acids Alleviate *In Vitro* Skeletal Muscle Dysfunction Induced by Oxidative Stress

**DOI:** 10.64898/2025.12.29.696779

**Authors:** Xinyue Lu, Olga L. Sharko, Vadim V. Shmanai, Mikhail S. Shchepinov, James F. Markworth

## Abstract

Excessive oxidative stress drives lipid peroxidation and contributes to skeletal muscle atrophy in a range of musculoskeletal diseases. Polyunsaturated fatty acids (PUFAs) are essential components of muscle cell membrane phospholipids and are especially susceptible to peroxidation due to the presence of double bonds. Currently, therapeutic options targeting lipid peroxidation to prevent muscle wasting are limited. Substituting the hydrogen atom at the bis-allylic position with deuterium could conceivably limit lipid peroxidation while retaining enzymatic PUFA metabolism. Here we investigated the potential role of deuterated PUFAs (D-PUFAs) in protecting against muscle cell dysfunction under conditions of elevated oxidative stress. Both native (H-) and deuterated (D-) forms of long chain PUFAs including arachidonic acid (ARA), eicosapentaenoic acid (EPA), docosapentaenoic acid (DPA), and docosahexaenoic acid (DHA) stimulated *in vitro* muscle cell growth and development in the absence of oxidative stress. D-ARA, D-EPA, D-DPA, and D-DHA each protected cultured myotubes against the deleterious effects of direct exposure to reactive oxygen species (ROS) by limiting lipid peroxidation. In contrast, H-ARA, H-EPA, H-DPA, and H-DHA each increased sensitivity to ROS-induced lipid peroxidation and exacerbated oxidative stress-induced muscle cell dysfunction. Deuterated short chain linoleic acid (D-LA), alpha linolenic acid (D-ALA), as well as D-ARA and D-EPA (but not D-DPA or D-DHA) also protected against the deleterious effects of ferroptosis inducer erastin on myogenic differentiation. Finally, D-PUFAs modulated local expression of endogenous antioxidant enzymes and muscle-specific protein ligases. Overall, our study suggests a promising role of D-PUFAs as novel therapeutics to protect against skeletal muscle dysfunction induced by oxidative stress.

## INTRODUCTION

Skeletal muscle is one of the largest components of the human body and is essential in mediating physical performance and tissue metabolism (1). Skeletal muscle atrophy refers to the involuntary loss of muscle mass, strength, and function, which could be caused by a wide range of pathophysiological conditions, including long-term muscle disuse, metabolic diseases (e.g., diabetes), muscle injuries, inflammatory diseases, cancer, and aging (2). In many diseases, prolonged muscle atrophy impairs physical function and worsens the underlying conditions, negatively impacting the prognosis and the life quality of patients.

Oxidative stress, characterized by an imbalance between the production of reactive oxygen species (ROS) and endogenous antioxidant defenses, plays a crucial role in modulating skeletal muscle health across various disorders (3). Muscle cells are regularly exposed to ROS produced by NADPH oxidase and the electron transport chain (ETC) in mitochondria during oxidative metabolism (4). During muscle regeneration, the production of ROS by M2 macrophages and neutrophils is required to facilitate phagocytosis and muscle regeneration, which can typically be maintained at the physiological level by endogenous antioxidant mechanisms (5). Nevertheless, an imbalanced redox state and excessive ROS accumulation in muscle fibers could lead to cellular damage, mitochondrial dysfunction, chronic inflammation, and muscle atrophy (6). For example, excessive ROS contributes to sarcopenia by activating muscle cell apoptosis, inducing neuromuscular junction degeneration, and inhibiting muscle regeneration capacity (7–10).

Ferroptosis is an iron-dependent dysregulation of the redox state, leading to non-apoptotic cell death (11). During ferroptosis, excessive ferrous iron (Fe^2+^) could participate in the Fenton reaction and generate oxidative radicals such as hydroxyl radical (·OH), which then primarily targets phospholipids located in plasma and mitochondrial membranes (12). Once initiated by free radical attack, lipid peroxidation could cause persistent membrane damage and induce ROS-dependent cell lysis (13, 14). As an important component of cellular membranes, polyunsaturated fatty acids (PUFAs) are especially susceptible to lipid peroxidation due to the presence of double bonds (15). The skeletal muscle cell membrane (sarcolemma) contains multiple omega-3 (n-3) and omega-6 (n-6) PUFAs, including short chain linoleic acid (LA, C18:2 n-6) and alpha linolenic acid (ALA, C18:3 n-3), and long chain arachidonic acid (ARA, C20:4 n-6), eicosapentaenoic acid (EPA, C20:5 n-3), and docosahexaenoic acid (DHA, C22:6 n-3) (16). Free radicals attacking PUFAs trigger a self-propagating chain reaction, which gives rise to lipid peroxides and secondary peroxidation products such as phospholipid hydroperoxide (PLOOH), malondialdehyde (MDA), and 4-hydroxynonenal (4-HNE), driving ferroptotic cell death and muscle atrophy (17, 18). In skeletal muscles, the accumulation of cytotoxic lipid peroxides disturbs mitochondrial homeostasis and enhances protein degradation, impairing myogenic differentiation and regeneration (19–24).

Under normal and physiological circumstances, ferroptosis can be prevented by an endogenous antioxidant mechanism involving glutathione peroxidase (GPX), which is activated by the Nrf2 pathway upon oxidative stress (25–27). Cystine enters the cell through the cysteine/glutamate antiporter System X_c_^-^ and is converted to glutathione (GSH), the essential substrate of GPX (28). Lipid peroxides can be reduced to lipid alcohols by GPX and thus terminate the chain reaction, which is coupled by the oxidation of GSH to glutathione disulfide (GSSG) (28, 29). However, this antioxidant defense can be disrupted by a deficiency of GSH and selenium as well as experimental compounds and pharmacotherapeutics (30). For instance, the RAS-selective lethal (RSL) compound erastin and the anti-cancer drug sorafenib block the entrance of cysteine by targeting System X_c_^-^, limiting the synthesis of GSH (31). Similarly, buthionine sulfoximine (BSO) irreversibly arrests γ-glutamyl cysteine synthetase and directly halts the synthesis of GSH, which inhibits the activity of GPX (32–34). Previous research has shown that the ferroptosis-inducers erastin and BSO promote sarcopenia-like phenotype and inhibit muscle stem cell activity (35–37). Moreover, GPX-4 knockout in mouse models drives ferroptosis and leads to severe muscle atrophy and motor neuron degeneration, confirming the important role of endogenous antioxidant mechanisms in mediating ROS-related muscle wasting (38).

In the past few decades, multiple therapeutic agents have been studied to target ROS-related disorders. One potential approach involves the pharmacological administration of chemically modified PUFAs where the hydrogen atoms in the bis-allylic positions are replaced by deuterium atoms (D-PUFAs). The substitution of deuterium inhibits the abstraction of hydrogen by free radicals, thus limiting the autoxidation of native PUFAs (H-PUFAs) (39, 40). Considering the important role of PUFAs in constituting cellular membranes, D-PUFAs may have a promising role in preventing lipid peroxidation caused by membrane free radical attack while also preserving the enzymatic metabolism of essential fatty acids. Previous research has shown a promising role of D-PUFAs in targeting neurodegenerative diseases caused by oxidative stress (41). Rich in DHA and ARA, the brain may be especially prone to ROS-mediated lipid peroxidation. Indeed, oxidative stress induced by lipid peroxidation contributes to Alzheimer’s disease (AD) and Parkinson’s disease (PD) by impairing glucose metabolism, oxidizing critical enzymes, upregulating neural inflammation, and promoting mitochondrial dysfunction (42–44). In aldehyde dehydrogenase 2 (*Aldh2*) null mice exhibiting an AD-like phenotype, diets containing deuterated linoleic acid (D-LA) and deuterated α-linolenic acid (D-ALA) rescued cognitive impairment when compared to native PUFAs (H-PUFAs) (45). D-PUFAs have also been shown to improve cell viability and increase longevity in various neurodegeneration model (41, 46, 47). Moreover, D-PUFA could benefit atherosclerosis and nonalcoholic steatohepatitis by inhibiting lipid peroxidation and lowering oxidative stress (48, 49). Despite the preventive role of D-PUFAs in several neurological and cardiovascular disorders involving oxidative stress, to our knowledge, research on the role of D-PUFAs in oxidative stress-induced skeletal muscle dysfunction has been limited.

In the current study, we utilized the murine C2C12 skeletal muscle cell line to explore the potential effects of D-PUFA supplementation in models of oxidative stress-induced muscle cell dysfunction. We show here that the administration of various individual D-PUFAs can protect against the deleterious effects of hydrogen peroxide (H_2_O_2_) on cultured myotubes *in vitro* and reverse the inhibitory effects of the ferroptosis inducer erastin on myogenic differentiation. These data demonstrate a novel role of D-PUFA in protecting skeletal muscle cells from chemical-induced oxidative stress, revealing a potential role of D-PUFAs as novel therapeutics to target ROS-related muscle dysfunction.

## MATERIALS AND METHODS

### Cell Culture

The C2C12 murine skeletal muscle cell line (ATCC, CRL-1772) was cultured in Dulbecco’s modified Eagle medium (DMEM, Gibco, 11995-065) containing 10% fetal bovine serum (FBS, Corning, 35-010-CV) and antibiotics (penicillin 100 U/mL, streptomycin 100 μg/mL, Gibco, 15140-122) at 37 °C in humid air with 5% CO_2_. When reaching a confluency of 70%, C2C12 cells were passaged and seeded into 12-well culture plates in growth media at a cell density of 2.5 × 10^4^/cm^2^ and allowed to proliferate and crowd for 72 hours. To induce differentiation of C2C12 myoblasts, the growth media was replaced by differentiation media (DM), DMEM containing 2% horse serum (HS, Gibco 26050088) and antibiotics. In experiments using C2C12 myotubes beyond day 3 post differentiation, the cell media was replaced with fresh DM on day 3.

### ROS Exposure

To compare the effects of H-PUFAs and D-PUFAs on mature myotubes, C2C12 cells were differentiated for three days and then pretreated with a 25 μM dose of individual H-PUFAs or D-PUFAs (ARA, EPA, DHA, or DPA) for 24 hours. To drive an acute increase in oxidative stress, C2C12 myotubes were then exposed to H_2_O_2_ (2 mM) with fresh H-PUFAs or D-PUFAs (25 μM) for a further 48 hours. H_2_O_2_ was dissolved in 1 × phosphate-buffered saline (PBS, Gibco, 10010-023), and a solvent vehicle was added to the control cells to match the concentration of PBS. C2C12 cells from this experiment were used to assess lipid peroxidation and myotube morphology as described in the next sections.

### Assessment of Intracellular Lipid Peroxidation

To detect intracellular lipid peroxidation, BODIPY 581/591 C11 (Invitrogen, D3861) was used. Day 3 post-differentiation myotubes were pretreated with H-PUFAs or D-PUFAs for 24 hours and then exposed to 2 mM H_2_O_2_ for 48 hours as described above. Live C2C12 myotubes were washed twice with 1 × PBS and then incubated with 5 μM BODIPY 581/591 C11 in the dark at 37 °C for 30 minutes. Live cells were imaged immediately using an automated fluorescent microscope (Echo Revolution) operating in inverted configuration equipped with green (FITC) and red (Texas Red) filters. Following live cell imaging, myotubes were fixed in 4% PFA and stained for analysis of myotube morphology as described in later section.

### Ferroptosis Induction

To compare the effects of H-PUFAs and D-PUFAs on myogenesis in the presence of an inducer of ferroptosis, C2C12 myoblasts were plated at a cell density of 2.5 × 10^4^/cm^2^ and allowed to proliferate for 48 hours. The cells were then pretreated with a 25 μM dose of individual H-PUFAs or D-PUFAs prepared in growth media for 24 hours. After that, the myoblasts were induced to differentiate with DM containing erastin (10 μM, Cayman Chemical, 17754) and fresh H-PUFAs or D-PUFAs (25 μM) for 72 hours. To investigate the roles of H-PUFAs and D-PUFAs in ferroptosis post-differentiation, confluent C2C12 myoblasts were induced to differentiate for 48 hours and then pretreated with a 25 μM dose of individual H-PUFAs or D-PUFAs prepared in DM for 24 hours. Then, myotubes continued to differentiate in the presence of erastin (10 μM) and fresh H-PUFAs or D-PUFAs (25 μM) for 72 hours. In all the experiments, erastin was dissolved in molecular grade ethanol (Fisher Scientific, BP2818-500) and a solvent vehicle was added to control cells to match the concentration of ethanol.

### H-PUFA and D-PUFA Treatment

D-PUFAs were provided by Dr. Mikhail S. Shchepinov in the form of free fatty acids. Deuterium-incorporated n-3 and n-6 PUFAs including D-LA, D-ALA, D-ARA, D-EPA, D-DPA, and D-DHA were tested for their ability to alleviate ROS-induced muscle cell atrophy (**Table 1**). As a comparison, C2C12 cells were supplemented with matching H-PUFAs including H-LA (Cayman Chemical, 90150), H-ARA (Cayman Chemical, 90010), H-EPA (Cayman Chemical, 90110), H-DPA (Cayman Chemical, 90165), and H-DHA (Cayman Chemical, 90310). H-PUFAs and D-PUFAs were dissolved in pure molecular-grade ethanol and stored at -80°C under nitrogen gas until use.

**Table 1.**
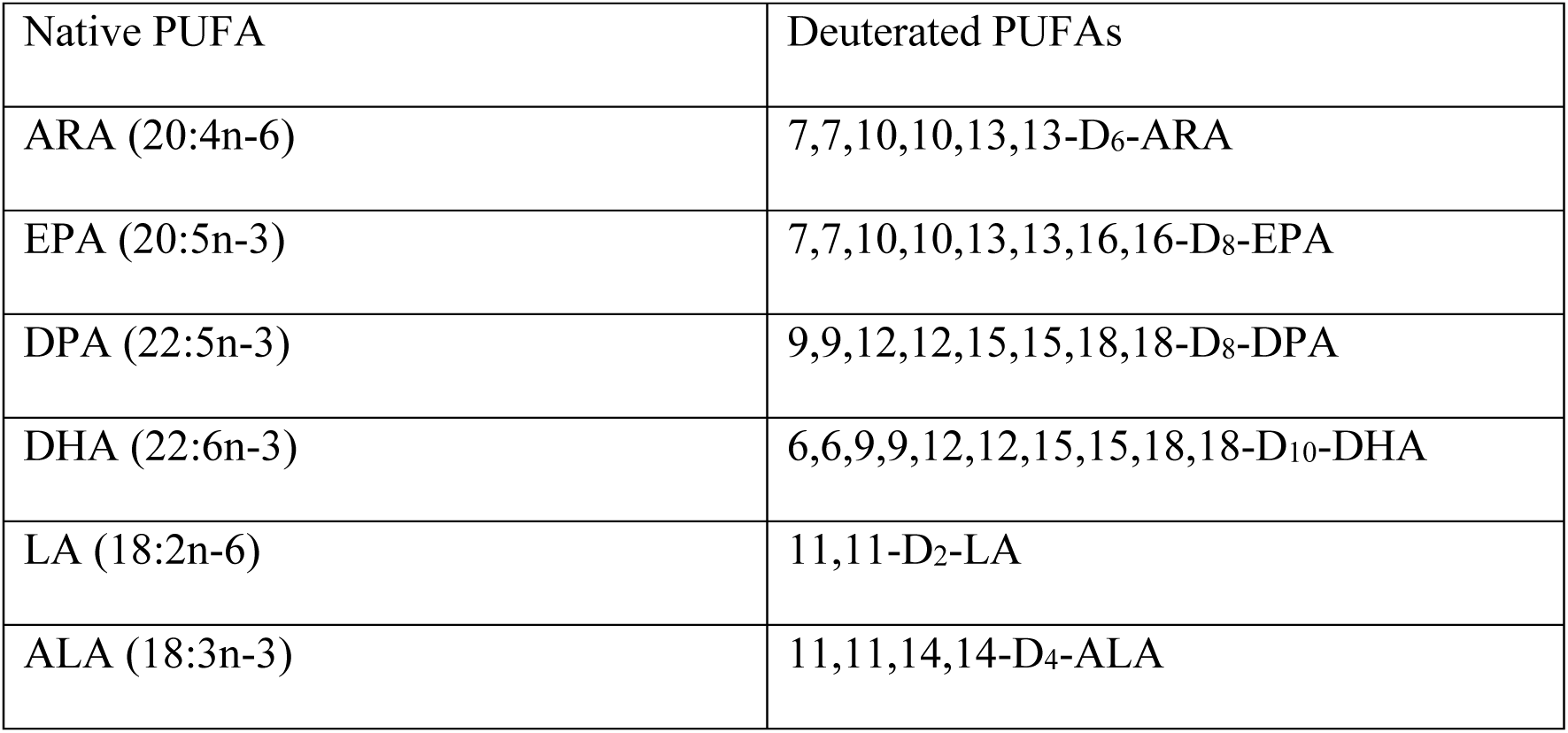
Native PUFAs and Enzymatically Converted Deuterated PUFAs Used in the Study.

### Immunocytochemistry

C2C12 myotubes were fixed in 4% paraformaldehyde (PFA, Electron Microscopy Sciences, 15710) for 30 minutes at 4°C and permeabilized with 0.1% Triton X-100 for 30 minutes at room temperature. The cells were then blocked in 1% bovine serum albumin (BSA, Sigma-Aldrich A3294-50G) for 1 hour at room temperature prior to overnight incubation at 4°C with primary antibodies against myogenin (F5Dc, DSHB, 1:100) and the sarcomeric myosin (MF20c, DSHB, 1:100). The following day, cells were washed in PBS and then incubated for 1 hour at room temperature with a mixture of Alexa Fluor conjugated secondary antibodies including Goat Anti-Mouse IgG2b Alexa Fluor 647 (Invitrogen, Thermo Fisher Scientific A-21242, 1:500) and Goat Anti-Mouse IgG1 Alexa Fluor 568 (Invitrogen A-21124, Thermo Fisher Scientific, 1:500), together with DAPI (Invitrogen, Thermo Fisher Scientific D21490, 2 μg/mL) to counterstain cell nuclei. Cells were washed with 1 × PBS at least three times before imaging.

### Image Analysis

Stained C2C12 cells were visualized using an automated fluorescent microscope (Echo Revolution) operating in inverted configuration. To avoid any potential investigator bias nine images were automatically captured from the same predetermined positions in each culture well using a 10 × Plan Fluorite objective. Overall cell density, myotube area, average number of myonuclei per myotube, and myotube diameter and were analyzed using ImageJ/FIJI as quantitative indices for myotube growth and differentiation. To determine mean myotube diameter, the ten largest myosin-positive cells from each image were manually measured at their widest uniform point perpendicular to the length of the cell using the straight-line selection tool. For branching myotubes, each branch was measured as a single myotube, and the region where the branches converge was excluded from analysis. Total cell number (DAPI^+^ cells/mm^2^) and myosin-positive (myotube) area (μm^2^) per field of view were quantified using a custom automated in-house plugin for Image J/FIJI. The total number of myosin-positive cells with ≥2 myonuclei (myotubes) was manually counted from the same FOVs. To determine the average number of myonuclei per myotube, total number of DAPI^+^ nuclei located with myosin^+^ cytoplasm was divided by total number of myotubes in the same fields of view.

### RNA extraction and RT-qPCR

For gene expression experiments confluent C2C12 myoblasts were induced to differentiate for 72 hours and then treated with a 25 μM dose of individual D-PUFAs in combination with 10 μM erastin for a further 24 hours. RNA was extracted using TRIzol reagent and RNA yield was determined using a NanoDrop 1000 UV/Vis Spectrophotometer (NanoDrop Technologies, E112352). After removing contaminating genomic DNA using Ambion^TM^ DNase I (Invitrogen, Thermo Fisher Scientific, AM2222), RNA (2 μg) was reverse-transcribed to cDNA using High-Capacity RNA-to-cDNA Kit (Thermo Fisher Scientific 4387406). RT-qPCR was performed in duplicate 10 μL reactions of PowerUp SYBR^TM^ Green Master Mix (Thermo Fisher Scientific, A25742) with 1 μM forward and 1 μM reverse primer (**Table 2**) on a 384-well QuantStudio^TM^ 5 Real-Time PCR Instrument (Thermo Fisher Scientific, A28135). Relative mRNA expression was calculated using the 2^-ΔΔCT^ method, with TATA-box binding protein (*Tbp*) used as the endogenous control.

**Table 2.**
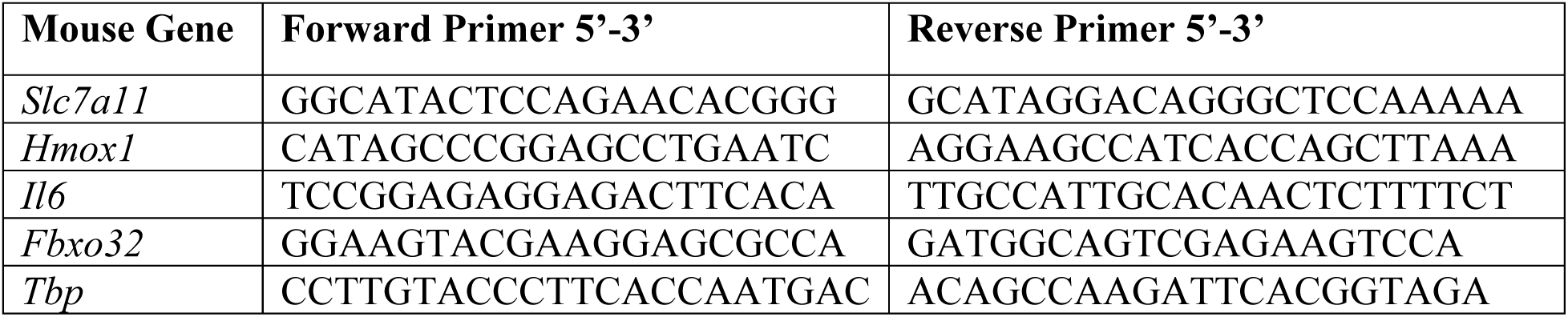
Primer Sequences for Genes used in RT-qPCR.

### Statistics

Statistical analysis was performed in GraphPad Prism 10. Data is shown as the mean ± SEM and raw data of three independent culture wells (considered to be biological replicates). Differences between treatment groups (1 factor with ≥3 levels) were detected by one-way analysis of variance (ANOVA) or two-way ANOVA (≥2 factors with ≥2 levels) followed by pairwise Holm-Šidák post-hoc tests. P ≤ 0.05 was used to determine statistical significance.

## RESULTS

### Both H-PUFAs and D-PUFAs Promote Myotube Fusion in the Absence of Oxidative Stress

Prior research has generated inconsistent findings regarding the potential impact of H-PUFAs upon skeletal muscle cell growth and development under normal cell culture conditions (50–53). Moreover, the potential for differential influences of H-PUFAs and D-PUFAs on skeletal muscle cells *in-vitro* have not been investigated previously. To determine the baseline effects of H-PUFAs and D-PUFAs on myogenesis in the absence of oxidative stress, confluent C2C12 myoblasts were induced to differentiate in the presence or absence of individual H-PUFAs or D-PUFAs for 72 hours (**Figure 1A**). Supplementation with pure individual H-PUFAs including H-ARA, H-EPA, H-DPA, and H-DHA did not influence overall muscle cell density (**Figure 1B**). Nevertheless, H-ARA, H-EPA, H-DPA, and H-DHA, each significantly increased mean myotube diameter (**Figure 1C**) and the average number of myonuclei per myotube **(Figure 1D).** Supplementation with pure individual D-PUFAs including D-ARA, D-EPA, D-DPA, and D-DHA also each increased myotube diameter (**Figure 1C**) and the average number of myonuclei per myotube (**Figure 1D**). Like H-PUFAs, corresponding D-PUFAs did not have any effect on total cell number except for D-ARA that slightly increased overall cell density (**Figure 1B**).

**Figure 1.**
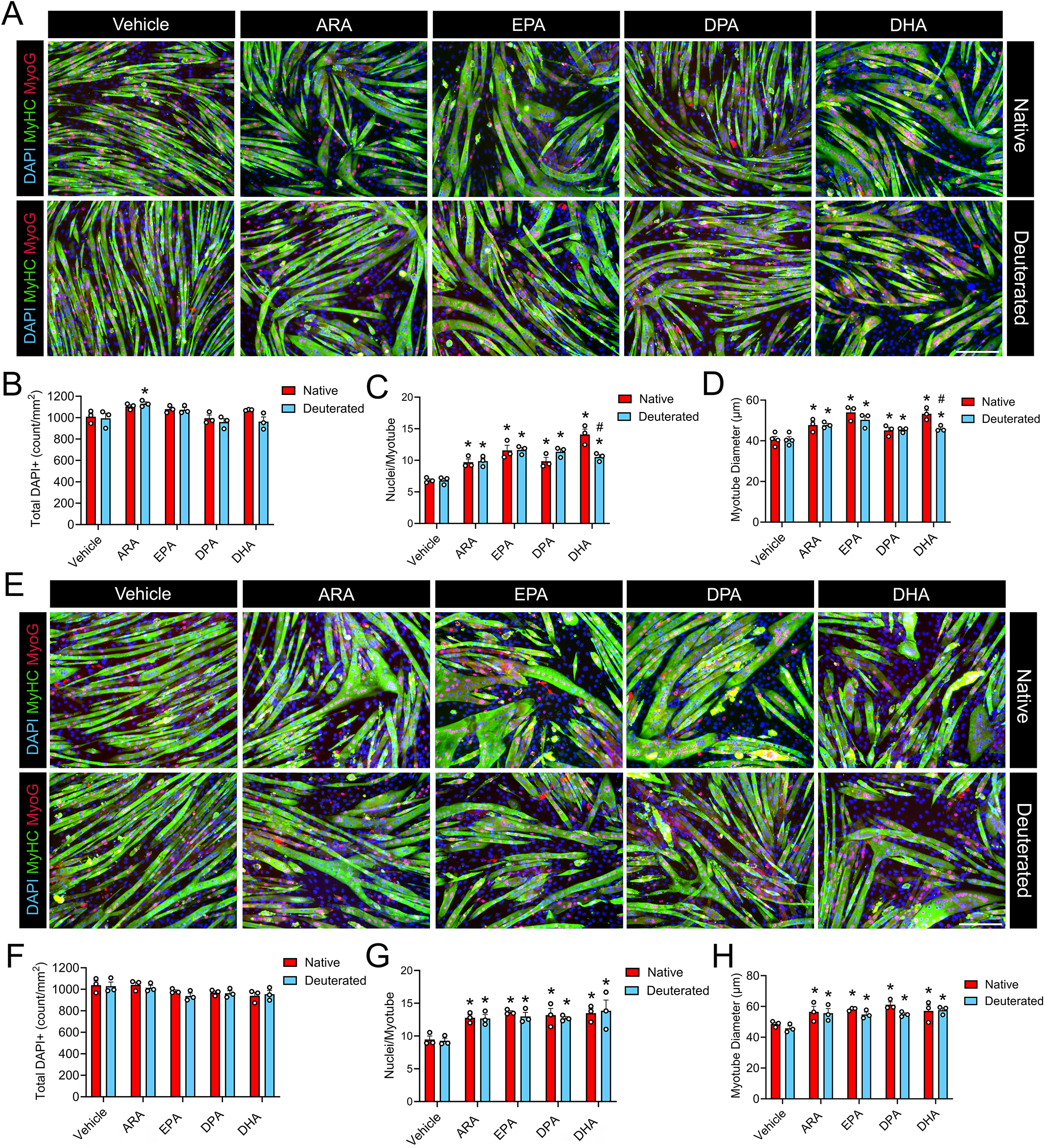
Both H-PUFAs and D-PUFAs increase myotube diameter in differentiating and mature myotubes in the absence of oxidative stress. **(A):** To assess the effect of H-PUFAs and D-PUFAs during differentiation, confluent C2C12 myoblasts were induced to differentiate for 72 hours in DM supplemented with a 25 µM dose of individual H-PUFAs or D-PUFAs. **(B-D):** Total DAPI^+^ cell count **(B)**, average myonuclei per myotube **(C)**, and average myotube diameter during differentiation **(D)** were quantified using ImageJ. **(E):** To assess the effect of H-PUFAs and D-PUFAs on mature myotubes, confluent C2C12 myoblasts were induced to differentiate for 72 hours in DM prior to supplementation with a 25 µM dose of individual H-PUFAs or D-PUFAs for 72 hours. **(F-H):** Total DAPI^+^ cell count **(F)**, average myonuclei per myotube **(G)**, and myotube diameter **(H)** were quantified using ImageJ. For both experiments, myotubes were fixed by 4% PFA and visualized after immunofluorescent staining for sarcomeric myosin (MF20c, colored green) and myogenin (F5Dc, colored red). Cell nuclei were counterstained with DAPI (blue). Scale bar is 200 µm. *P<0.05 for difference compared to vehicle, #P<0.05 for difference between H-PUFA and D-PUFA.

To determine the influence of H-PUFAs and D-PUFAs on mature myotubes in the absence of oxidative stress, day 3 post-differentiation C2C12 myotubes were treated with either H-PUFAs or D-PUFAs (25 μM) for 72 hours (**Figure 1E**). While no change was detected in total cell number (**Figure 1F**), both individual H-PUFAs and D-PUFAs significantly increased myotube diameter (**Figure 1G**) and the average number of myonuclei per myotube (**Figure 1H**). These results suggest that both native and deuterated long chain PUFAs can induce hypertrophy and myonuclear accretion in healthy C2C12 myotubes under standard cell culture conditions. Therefore, anabolic effects of H-PUFAs upon muscle cells do not appear to be impacted positively or negatively by deuterium reinforcement.

### Deuterium Incorporation Alleviates the Deleterious Effect of H-PUFAs on Myotubes Exposed to Oxidative Stress by Decreasing Intracellular Lipid Peroxidation

To test whether native and deuterated PUFAs might have divergent effects under conditions of high oxidative stress, day 3 post-differentiation C2C12 myotubes were pretreated with either H-PUFAs or D-PUFAs for 24 hours. Then the myotubes were exposed to H_2_O_2_ (2 mM) for a further 48 hours in the continued presence of respective PUFA substrates (**Figure 2A**). H_2_O_2_ treatment induced marked cell death as indicated by an overall reduction in muscle cell density (**Figure 2B**), myotube area (**Figure 2C**), and the average number of myonuclei per myotube (**Figure 2D**). Supplementation with H-ARA, H-EPA, H-DHA, and H-DPA each greatly exacerbated these deleterious effects of H_2_O_2_ treatment, leading to marked reductions in total cell number (**Figure 2B**), myotube area (**Figure 2C**), and the average number of myonuclei per myotube (**Figure 2D**). Nevertheless, these negative effects of H-PUFAs on myotubes exposed to oxidative stress were not observed in response to D-PUFA treatments (**Figure 2A**). Rather, when compared to corresponding H-PUFAs, supplementation with each individual D-PUFA increased overall muscle cell density (**Figure 2B**), myotube area (**Figure 2C**), and nuclei/myotube (**Figure 2D**). Moreover, D-PUFA supplementation greatly increased the average number of myonuclei per myotube when compared to H_2_O_2_ alone (**Figure 2D**). We also used the lipid peroxidation sensor BODIPY™ 581/591 C11 to visualize oxidized intracellular lipids. BODIPY™ 581/591 C11 is incorporated into cell membrane phospholipids and shifts from red to green fluorescence emission upon lipid peroxidation. Mature C2C12 myotubes exposed to H_2_O_2_ exhibited more oxidized lipids, and this response was further increased by H-ARA treatment (**Figure 2E**). On the other hand, myotubes receiving D-ARA treatment generated less oxidized lipids (**Figure 2E**). As expected, myotubes treated with H-EPA, H-DPA, or H-DHA showed even higher levels of lipid peroxidation than H-ARA in myotubes subsequently exposed to H_2_O_2_, presumably due to the higher number of double bonds and sensitivity to free radical attack. On the other hand, deuterium substitution limited the oxidation of EPA, DPA, and DHA in the presence of H_2_O_2_ (**Figure 2E**).

**Figure 2.**
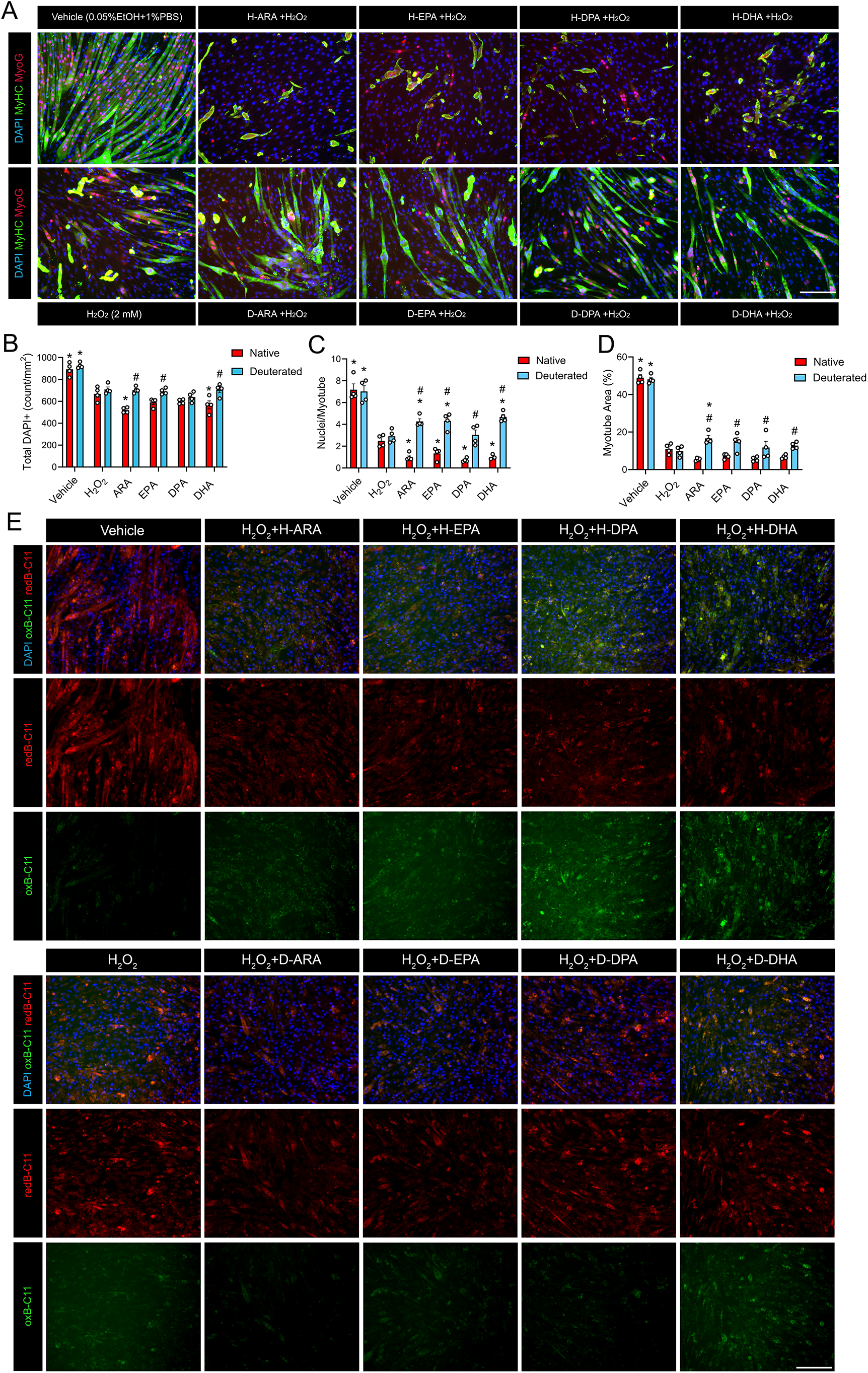
Deuteration protects skeletal muscle cells against the deleterious effects of H-PUFAs when exposed to H_2_O_2_. (**A):** To assess the effect of H_2_O_2_ on mature myotubes, confluent C2C12 myoblasts were induced to differentiate for 48 h in DM and then pre-treated with a 25 µM dose of individual H-PUFA or D-PUFA for a further 24 h. Subsquently, myotubes continued to grow in fresh DM with 2 mM H_2_O_2_ and a fresh 25 µM of H-PUFAs or D-PUFAs for 48 hours. Myotubes were fixed in 4% PFA and visualized after immunofluorescent staining for sarcomeric myosin (MF20c, colored green) and myogenin (F5Dc, colored red). Cell nuclei were counterstained with DAPI (blue). Scale bar is 200 µm. **(B-D):** Total DAPI^+^ cell count (**B**), average myonuclei per myotube (**C**), and myotube area (**D**) were quantified using ImageJ. (**E):** Bodipy C11 staining of C2C12 cells receiving H-PUFA or D-PUFA in the presence of H_2_O_2_. Oxidized lipids are in green and reduced lipids are in red. *P<0.05 for difference of PUFAs compared to H_2_O_2_, #P<0.05 for difference between H-PUFA and D-PUFA.

### Differential Effects of D-PUFAs on Myoblasts and Myotubes Receiving Erastin

To test the impact of ferroptosis-induced lipid peroxidation on myogenesis and whether deuterium incorporated PUFAs might potentially protect differentiating muscle cells against ferroptosis, we pretreated proliferating C2C12 myoblasts with individual H-PUFAs or D-PUFAs for 24 hours. Cells were then induced to differentiate in the presence of the ferroptosis-inducer erastin in the continued presence of H- or D-PUFAs for 72 hours (**Figure 3A**). Treatment with erastin alone at the onset of myogenic differentiation did not impact overall muscle cell density (**Figure 3B**) but did significantly impair myotube diameter (**Figure 3C**) and myonuclear accretion as indicated by a lower average number of myonuclei per myotube (**Figure 3D**). Treatment with H-ARA, H-EPA, H-DPA, and H-DHA during myogenic differentiation each markedly exacerbated the deleterious effects of erastin, resulting in complete cell death (**Figure 3B**). In contrast, treatment with D-PUFAs including D-ARA, D-EPA, D-DPA, or D-DHA did not result in such cell death (**Figure 3B**). In fact, D-ARA and D-EPA significantly increased both the average diameter of developing myotubes (**Figure 3C**) and the average number of myonuclei per myotube (**Figure 3D**). These apparent protective effects of D-PUFAs were not observed in cells receiving D-DPA or D-DHA (**Figure 3B**).

**Figure 3.**
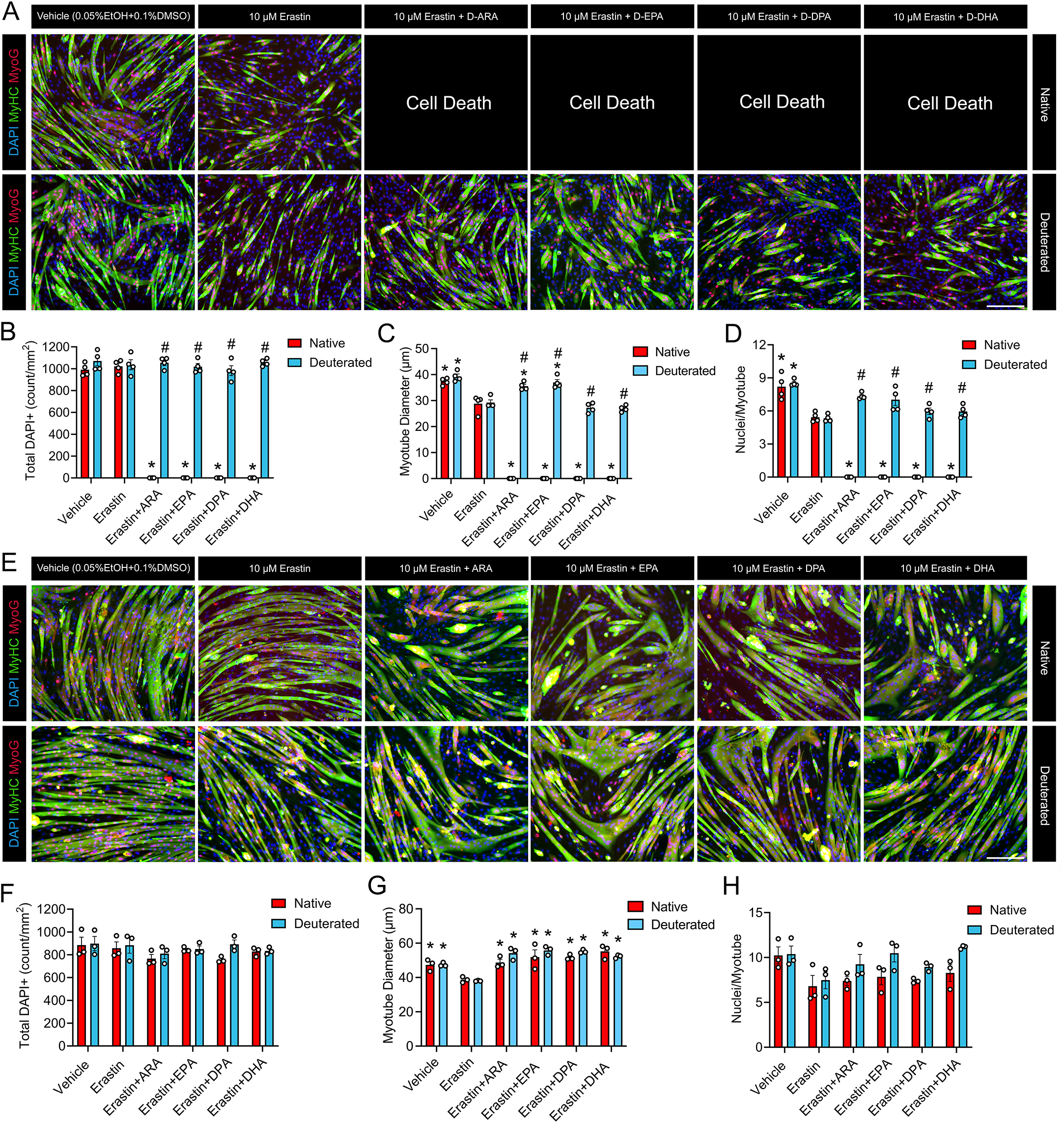
Individual D-PUFAs show differential effects on C2C12 exposed to erastin during and post differentiation. **(A)** To assess the effect of erastin during differentiation, proliferating C2C12 myoblasts were pretreated with a 25 µM dose of individual D-PUFAs including D-ARA, D-EPA, D-DPA, or D-DHA for 24 hours. Confluent myoblasts were then induced to differentiate in DM containing ferroptosis inducer erastin (10 µM) and fresh D-PUFAs (25 µM) for 72 hours. **(B-D):** Total DAPI^+^ cell count (**B**), average myonuclei per myotube (**C**), and average myotube diameter (**D**) were measured using ImageJ. **(E):** To assess the effect of erastin on mature myotubes, confluent C2C12 cells were induced to differentiate in DM for 48 hours and then pretreated with individual D-PUFAs for 24 hours. Subsquently, myotubes continued to grow in fresh DM containing erastin (10 µM) and fresh D-PUFAs for 72 hours. **(F-H):** Total DAPI^+^ cell count (**F**), average myonuclei per myotube (**G**), and myotube diameter (**H**) were quantified using ImageJ. Myotubes were fixed and visualized after immunofluorescent staining for sarcomeric myosin (MF20c, colored green) and myogenin (F5Dc, colored red). Cell nuclei were counterstained with DAPI (blue). Scale bar is 200 µm. *P<0.05 for difference during differentiation, #P<0.05 for difference post differentiation.

To test the effect of erastin on post-mitotic muscle cells, day 2 post-differentiation myotubes were pre-treated with H- or D-PUFAs for 24 hours prior to addition of erastin for a further 72 hours in the continued presence of respective PUFA substrates. When administered to mature myotubes, erastin clearly had less obvious detrimental effects on post-mitotic myotubes when compared to differentiating myoblasts (**Figure 3E**). Nevertheless, erastin treatment did still significantly reduce average myotube diameter (**Figure 3G)**. Furthermore, supplementation with D-ARA, D-EPA, D-DPA, and D-DHA each protected against this deleterious effect of erastin on myotube diameter (**Figure 3G**). In mature myotubes receiving erastin, the deleterious effect of H-PUFAs seen during differentiation were not observed (**Figure 3E**). Rather H-ARA, H-EPA, H-DPA, and H-DHA each protected against the deleterious effects of erastin on myotube diameter to a comparable extent to corresponding D-PUFA (**Figure 3G**). Erastin, H-PUFAs, and/or D-PUFAs did not have any significant effect on the average number of myonuclei per myotube when treatments were administered post-differentiation (**Figure 3H**).

### Protective Effects of D-PUFAs against Ferroptosis on Myogenic Differentiation are Dose-dependent

We next aimed to study whether the rescuing effects of D-PUFAs against ferroptosis induced muscle cell dysfunction are dose-dependent. D-ARA was chosen as a representative of long-chain D-PUFAs due to its potent protection against both H_2_O_2_ and erastin (see **Figures 1-3**). We also sought to test whether deuterium reinforced short-chain PUFAs including ALA (C18:3 n-3) and LA (C18:2 n-6) might have similar protective effects on skeletal muscle cells since they are anticipated to be enzymatically converted to longer chain PUFAs while preserving the deuterium. Proliferating C2C12 myoblasts were pre-treated with increasing doses of D-ARA, D-LA, or D-ALA for 24 h prior to induction of myogenic differentiation in the presence of erastin for 72 hours (**Figure 4A**). Consistent with earlier experiments, erastin greatly reduced myotube diameter (**Figure 4B**) and the average number of myonuclei per myotube (**Figure 4C**). Pretreating with D-ARA at a dose of 25 μM and 50 μM significantly reversed the deleterious effects of erasin on myotube diameter (**Figure 4B**) and nuclei/myotube (**Figure 4C**). However, at a 100 μM dose, the benefits of D-ARA were no longer observed for either myotube diameter (**Figure 4B**) or the average nuclei per myotube (**Figure 4C**). Treatment with either ALA or LA at a dose of 25 μM and 50 μM (but not 100 μM) also significant reversed the deleterious effects of erastin on myotube diameter (**Figure 4D & Figure 4F**) and myonuclei/myotube (**Figure 4E** & **Figure 4G**). Staining of neutral lipid droplets by BODIPY 493/503 revealed that treatment of mature myotubes exposed to erastin with high doses of D-ARA, D-ALA, and D-LA each led to marked lipid droplet accumulation (**Figure 4H**). The loss of protective effects at high doses of D-ARA, D-ALA, and D-LA at high doses (e.g., ≥100 µM) could potentially be related to the excessive accumulation of deuterium reinforced lipids in the cell membrane.

**Figure 4.**
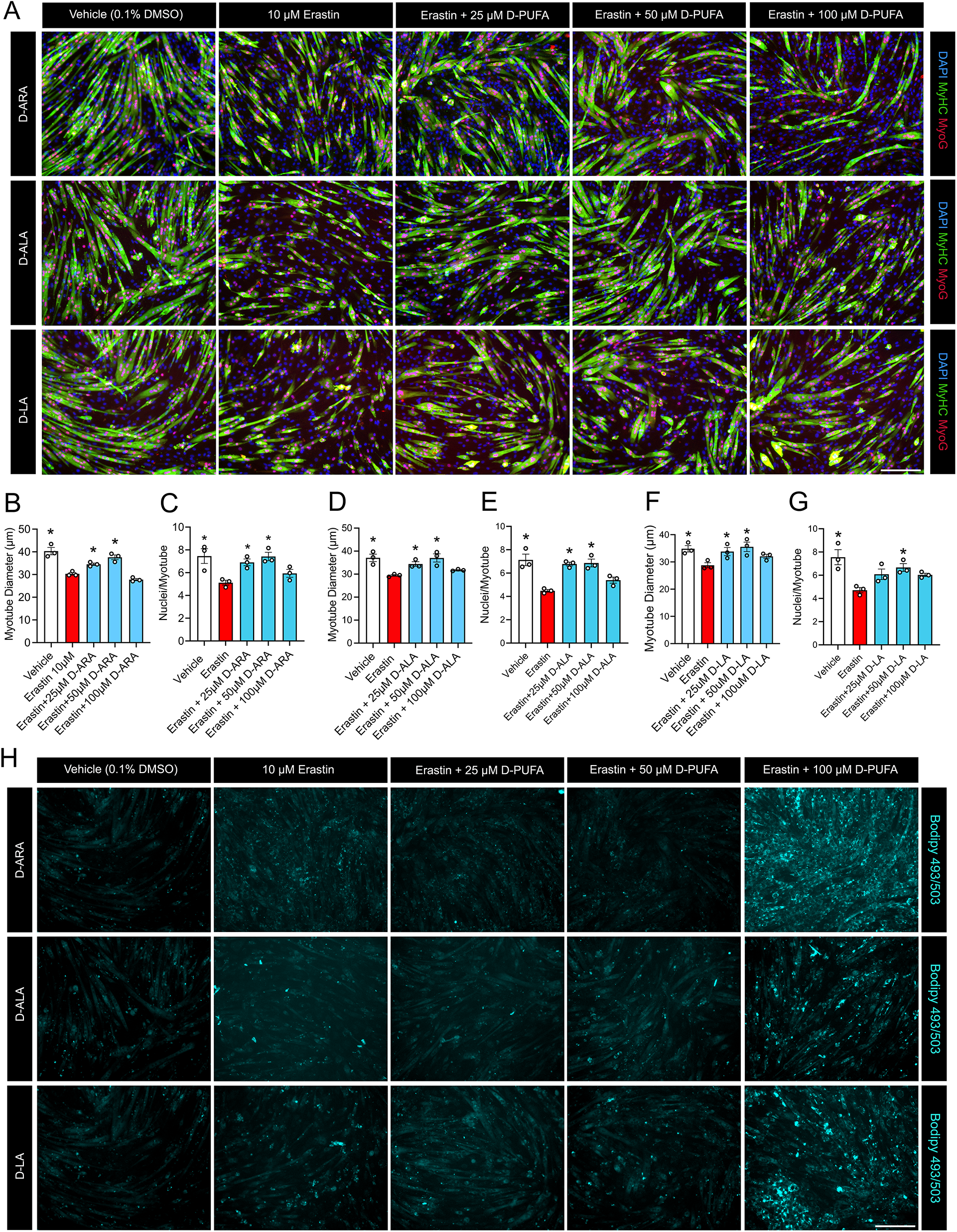
The effects of D-PUFAs on C2C12 exposed to erastin are dose dependent. **(A):** To assess the effect of D-PUFA dose during differentiation, proliferating C2C12 myoblasts were pretreated with an increasing dose of D-ARA, D-ALA, or D-LA (0, 25, 50, 100 µM) for 24 hours. Confluent myoblasts were then induced to differentiate for 72 h in DM containing erastin (10 µM) and fresh D-ARA, D-ALA, or D-LA at respective concentrations. Myotubes were fixed and visualized after immunofluorescent staining for sarcomeric myosin (MF20c, colored green) and myogenin (F5Dc, colored red). Cell nuclei were counterstained with DAPI (blue). Scale bar is 200 µm. **(B-G):** Myotube diameter (**B, D, & F**) and average nuclei per myotube (**C, E, & G**) was measured in cultures receiving treatment with D-ARA (**B-C**), D-ALA (**D-E**), and D-LA (**F-G**). **(H):** Neutral lipid stained by BODIPY 493/503 in the dose response experiment. Representative images shown in **(A)** and **(H**) were taken from the same fields of view. *P<0.05 for difference compared to cells receiving erastin alone.

### Erastin and D-PUFAs Regulate Genes Related to Antioxidant Responses in Mature Myotubes

To assess the underlying molecular mechanisms responsible for the protective effects of D-PUFAs on ferroptosis induced muscle cell dysfunction we measured the local expression of endogenous antioxidant genes. Day 3 post-differentiation myotubes were co-treated with erastin and individual D-PUFAs for 24 hours. Erastin markedly upregulated mRNA expression of Solute Carrier Family 7 Member 11 (*Slc7a11*) (**Figure 5A**), which could potentially be a compensatory mechanism to mitigate the malfunction of SLC7A11 protein (54, 55). In the absence of erastin D-PUFA supplementation did not influence the expression of *Slc7a11*. When stimulated by erastin, however, D-LA tended to reduce the *Slc7a11* expression (P=0.0537) while D-DHA rather further elevated *Slc7a11* expression (**Figure 5A**). As an intracellular source of iron, hemoxygenase-1 (HMOX-1, also known as HO-1) plays an indispensable role in activating erastin-induced ferroptosis (56). Erastin treatment significantly increased expression of *Hmox1* in C2C12 myotubes (**Figure 5B**). Co-treatment with D-LA suppressed the expression of *Hmox1* induced by erastin, whereas D-DHA rather increased *Hmox1* expression (**Figure 5B**). Erastin also increased the expression of the muscle-specific E3 ubiquitin ligase Atrogin-1 (*Fbxo32*), a well-established marker of skeletal muscle atrophy (57) (**Figure 5C**). Co-treatment with D-LA, D-DPA, and D-DHA each significantly downregulated erastin-induced *Fbxo32* expression (**Figure 5C**). Finally, erastin also significantly suppressed the expression of the cytokine interleukin 6 (*Il6*) (**Figure 5D**). D-ARA stimulated *Il6* expression irrespective of erastin treatment, but no significant effect was observed with other D-PUFAs (**Figure 5D**).

**Figure 5.**
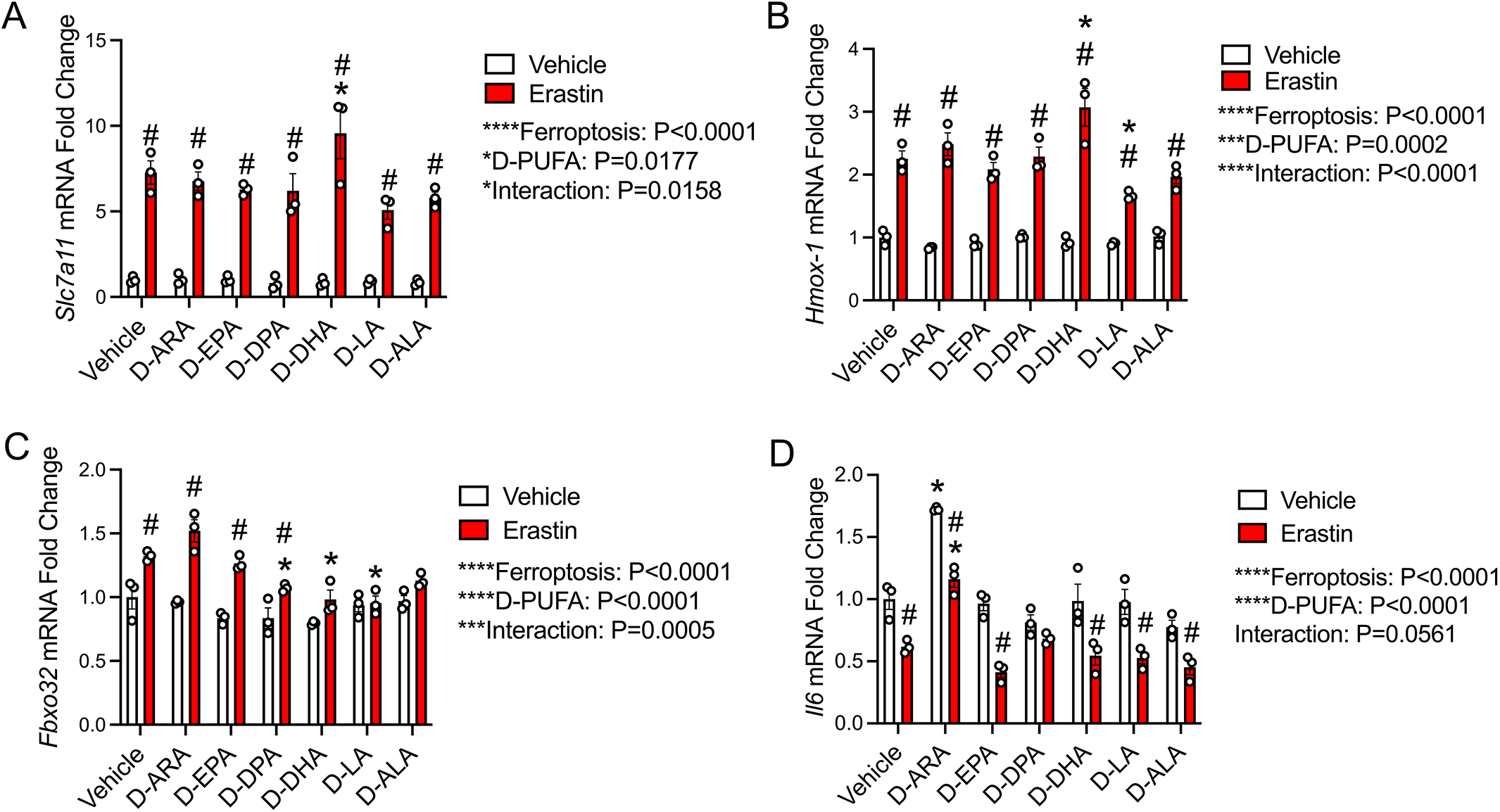
D-PUFAs regulate expression of endogenous antioxidant enzymes and ubiquitin ligases. At 3 days post-differentiation, mature C2C12 myotubes were co-treated with erastin (10 µM) and D-PUFAs (25 µM) in fresh DM for 24 hours. Cell mRNA expression of *Scl7a11* (**A**), *Hmox1* (**B**), *Fbxo32* (**C**), and *Il6* (**D**) were assessed using RT-qPCR as described in methods. Gene expression was normalized to *Tbp*. *P<0.05 for effect of D-PUFAs compared to vehicle, #P<0.05 for effect of erastin.

## DISCUSSION

Oxidative stress impedes myogenesis and contributes to skeletal muscle atrophy through different pathways, one of which is activating lipid peroxidation of PUFA species on the muscle cell membrane (19, 23, 58). It has been shown that oxidative stress negatively influences the development of skeletal muscles (6). Furthermore, previous research has demonstrated a protective role of deuterated PUFAs against muscle atrophy in a diabetic mouse model (59). However, no research has directly compared the effects of H-PUFAs and D-PUFAs on skeletal muscle cells under high oxidative stress. In this study, we investigated the differential roles of H-PUFAs and D-PUFAs in mediating skeletal muscle cell dysfunction induced by oxidative stress.

Previous research shows that acute increases in H_2_O_2_ level could result in a decrease in net muscle protein concentration and altered sarcomeric morphology (60). We found that loading mature myotubes with native PUFAs greatly exacerbated the deleterious effects of exposure to high levels of ROS (H_2_O_2_). On the contrary, the substitution of hydrogen atoms with deuterium in bis-allylic positions of PUFAs limited the deleterious effect of H_2_O_2_ on C2C12 myotubes. H_2_O_2_ produces ·OH through the Fenton reaction to initiate lipid peroxidation, potentially contributing to the deleterious effect of native PUFA species (61). Assessment of lipid peroxidation showed that the incorporation of native PUFAs to muscle cell membrane in the presence of H_2_O_2_ results in substantial accumulation of oxidized lipids, inhibiting myotube growth and inducing cell death. The amount of lipid peroxidation was proportional to the total number of bis-allylic groups in the PUFA substrate which is consistent with previous findings on the kinetics of PUFA autoxidation (62).

Erastin drives ferroptosis by inhibiting the entrance of cysteine and blocking the GSH-dependent antioxidant pathway (31). In differentiating C2C12 myoblasts, we observed that erastin-induced ferroptosis significantly blocked myotube fusion and suppressed the expression of myosin in accordance with prior findings (36). Moreover, the deleterious effect of ferroptosis was exacerbated by native PUFAs, inducing oxidative stress-related cell death. The detrimental effect of the ferroptosis inducer erastin on differentiating myoblasts was partially rescued by D-ARA, D-EPA but not D-DPA or D-DHA. Interestingly, a previous study showed that D-DHA exhibited a stronger protection against lipid peroxidation compared to D-ARA and D-EPA in an *in-vitro* liposome model composed of single phospholipids, and the required incorporation rate of D-DHA and D-ARA for effective protection against liposome ferroptosis (63, 64). The lower protective efficiency of D-DHA we observed in the skeletal muscle cell ferroptosis model could be related to the lipid profile in the muscle cell membrane and the differential incorporation rate of D-PUFAs in skeletal muscle cells. Previous research suggests that human skeletal muscle phospholipids contain 36.2% LA, 16.8% ARA, and 1.3% DHA, which could contribute to the higher protective efficiency of D-LA and D-ARA in C2C12 (65). Moreover, there could be a potential threshold in the protective effect of D-PUFAs on skeletal muscle cells exposed to oxidative stress. Extreme oxidative stress causing extensive damage to multiple cell compartments may not be fully resolved by the simple incorporation of deuterium into bilayer phospholipid membranes.

In addition, the protective effect on myotube differentiation against ferroptosis was also observed with short chain PUFAs including D-ALA and D-LA. ALA and LA are essential fatty acids in humans and play important roles in modulating lipid profile and metabolism (66). Dietary supplementation of ALA increases sarcolemma transport of lipids and changes skeletal muscle membrane lipid composition, increasing n-3 PUFAs and decreasing n-6 ARA (67). LA regulates muscle membrane fluidity and can be enzymatically converted to n-6 ARA, a key lipid component of muscle cell membrane (68). It has been shown that native LA drives ferroptosis in cancer cells when exposed to RSL-3, while D-LA suppresses lipid peroxidation (69). A previous study also shows that D-LA and D-ALA suppress F_2_-isoprostanes (F_2_-isoPs) and reduce prostaglandin F_2α_ (PGF_2α_) in brain tissues of mice with sporadic AD (45). Moreover, n-3 D-ALA and n-6 D-LA could be enzymatically converted to long chain D_4_-EPA and D_4_-ARA, correspondingly, thus allowing the extended incorporation of deuterium and protection against autoxidation (70). Our results show that D-ALA and D-LA protect against ferroptosis-induced inhibition on myotube differentiation, which could be related to the incorporation of deuterium into longer chain PUFAs during enzymatic metabolism.

Moreover, we found that the rescuing effect of D-PUFAs was dose-dependent. When administered at does below 50 µM, we observed a protective effect of D-ARA, D-ALA, and D-LA on myogenic differentiation during ferroptosis. However, when administered at a dose of 100 µM to differentiating myoblasts, D-ARA, D-ALA, and D-LA did not exhibit any protective effects on myotube formation, which is associated with lipid droplet accumulation. Although it has been shown that the formation of lipid droplets promotes myogenesis and fusion of myoblasts by driving microfilament remodeling, excessive accumulation of lipid droplets could repress energy production and induce higher levels (71, 72). Additionally, erastin enhanced the accumulation of lipid droplets without any D-PUFA substrates, suggesting a potential crosstalk between ferroptosis and dysregulated lipid metabolism, which could contribute to the inhibitory effect of erastin on myotube formation and fusion.

We did observe a difference in erastin tolerance between C2C12 myoblasts and myotubes. While erastin-induced ferroptosis substantially inhibited myogenesis and differentiation, mature C2C12 myotubes seemed to be more tolerant to ferroptosis, which could be related to the difference in mitochondrial metabolism, cellular stress, and cell-death resistance (73, 74). It has been shown that differentiated myotubes have decreased expression of poly(ADP-ribose) polymerase 1 (PARP-1), leading to more preserved mitochondrial function and lower sensitivity to cytotoxicity induced by oxidative stress (73). Meanwhile, mature myotubes exhibit a shift of mitochondrial metabolism to a higher level of oxidative phosphorylation and increased respiratory capacity (64, 73). Future research could potentially investigate the molecular mechanisms by which erastin differentially interacts with PUFAs in differentiating myoblasts when compared to post-mitotic myotubes.

We also found a regulatory role of D-PUFAs on endogenous antioxidant gene expression in response to oxidative stress. Erastin markedly induced mRNA expression of the cystine/glutamate transporter *Scl7a11*, an essential component of the GPX4-dependent antioxidant mechanism. Previous studies show that erastin induces ferroptosis by allosterically inhibiting the protein activity of SLC7A11, the functional subunit of cystine/glutamate antiporter System X_c_^-^, which is essential for the antioxidant defense dependent on GSH metabolism (29, 75, 76). The increased mRNA level of *Slc7a11* could be a compensatory response induced by repressed cystine uptake to restore redox balance under oxidative stress (77, 78). However, such compensation could also lead to a higher demand for NADPH and glucose metabolism, potentially leading to a higher vulnerability to redox imbalance and consequently disulfidptosis (79, 80). Our results show that the administration of D-LA tends to downregulate the expression of *Scl7a11* in C2C12 myotubes and alleviate oxidative stress, suggesting D-LA may alleviate ferroptosis by mediating cystine uptake and endogenous antioxidant mechanisms in addition to directly limiting autoxidation of LA. On the other hand, D-ARA, D-EPA, and D-ALA did not show any influence on *Slc7a11* expression, indicating they protect myotubes from ferroptosis through other mechanisms.

Consistent with prior research, we found that *Hmox1* expression was elevated during erastin-induced ferroptosis (81). HMOX1 plays a complex role in mediating oxidative stress, eliciting both protective and cytotoxic effects (82). Activation of HMOX1 protects cancer cells against H_2_O_2_-induced oxidative stress by upregulating ferritin synthesis and decreasing redox active iron (83). However, excessive HMOX1 is rather related to the accumulation of reactive ferrous and could lead to non-canonical ferroptosis (84, 85). Similarly, in human fibrosarcoma cells, the activation of HMOX-1 facilitates the accumulation of lipid peroxidation under oxidative stress, which further sensitizes the cells to erastin-induced ferroptosis (56). Moreover, overexpression of *Hmox1* contributes to elevated ER stress and disrupted mitochondrial homeostasis (86). In this study, the increased expression of *Hmox1* was attenuated by D-LA but further elevated by D-DHA, which is in line with their effects on *Slc7a11* expression. Nevertheless, due to the scope of this paper, we did not determine the intracellular distribution of iron and the crosstalk between D-PUFAs and iron status during ferroptosis. Future studies could identify the interaction between D-PUFAs, HMOX1, and iron regulation during erastin-induced ferroptosis.

We also detected a decrease in *Il-6* expression caused by erastin, which was elevated by D-ARA. IL-6 plays complicated roles in regulating muscle inflammation, differentiation, and regeneration (87). Muscle-derived IL-6 is essential for myonuclear accretion and satellite cell-induced muscle hypertrophy (88). As a myokine, IL-6 has been shown to be crucial during muscle differentiation and muscle regeneration (89). The suppressed expression of IL-6 by erastin could be related to the blunted myotube fusion and consequently reduced myotube diameter we observed. D-ARA upregulated the expression of *Il-6* both with and without erastin, which could potentially contribute to its benefit on myotube fusion.

In addition, our gene data shows that D-DHA, D-ALA, and D-LA downregulated erastin-induced mRNA expression of Atrogin-1 (*Fbxo32*), indicating their potential role in mitigating the activation of the muscle-specific ubiquitin ligase pathway induced by ferroptosis. Crosstalk between oxidative stress and muscle protein turnover has been demonstrated in multiple muscle wasting models (90–92). In cancer-induced skeletal muscle wasting (cachexia), oxidative stress interacts with tumor-secreted inflammatory factors and enhances muscle protein degradation by activating the nuclear factor kappa B (NF-κB) pathway (3, 7–10, 93, 94). Moreover, the chemotherapeutic agent cisplatin has been found to induce intracellular ROS production and mitochondrial dysfunction, which could exacerbate muscle atrophy by activating the forkhead box O (FOXO)-Atrogin*/*MuRF1 pathway (95, 96). It has been shown that the supplementation of exogenous antioxidants remarkably alleviates protein degradation by repressing MuRF1 and preventing the downregulation of MyoD and myogenin (97). Similarly, in sarcopenia, oxidative stress contributes to the activation of FOXO1 and FOXO3, driving the muscle protein ligase pathway and autophagic activity (98–100). Previously, research demonstrated that H-DHA protected against palmitate-induced muscle wasting by suppressing FOXO3*/*Atrogin-1 and activating the protein kinase B (Akt) pathway (101). The apparent role of D-DHA in regulating Atrogin-1 expression suggests a protective effect of n-3 DHA on muscle catabolism irrespective of deuterium substitution. However, we did not observe any effect of D-EPA on Atrogin-1 mRNA expression in erastin treated myotubes despite its protective effect against ferroptosis related muscle cell wasting.

One limitation of the current study is the lack of an *in vivo* study to investigate the potential role of D-PUFAs in skeletal muscle function with or without oxidative stress. Future research in animal models could focus on the potential influence of dietary D-PUFA supplementation on skeletal muscle wasting induced by oxidative stress. The influences of D-PUFAs on skeletal muscle protein synthesis, autophagic/lysosomal activity, and myonuclear apoptosis were also not determined in the current study. As a result, one future direction is to investigate other potential mechanisms where D-PUFAs regulate oxidative stress in skeletal muscles. Moreover, future research could determine the incorporation rate or oxidation kinetics of H-PUFAs and D-PUFAs under different stressors.

## CONCLUSIONS

The current study provides evidence that deuterium-incorporated PUFAs protect skeletal muscle cells against *in-vitro* exposure to elevated oxidative stress, one of the key factors involved in the pathophysiology of sarcopenia and cancer cachexia. Specifically, we show that unlike H-PUFAs that sensitize muscle cells to the deleterious effects of oxidative stress, D-PUFAs can prevent the negative effects of ferroptosis-induces on myogenesis. The beneficial effects of D-PUFAs on muscle cell growth and development under conditioned of high oxidative stress appear to be related to their ability to limit lipid peroxidation.

## ACKNOWLEDGEMENTS

The authors thank Shihuan Kuang (Duke University, formerly of Purdue University) for laboratory support and helpful discussions. The MF20 (developed by Fischman, D.A.) and F5D (developed by Wright, W.E.) monoclonal antibodies were obtained from the Developmental Studies Hybridoma Bank (DSHB), created by the NICHD of the NIH and maintained at The University of Iowa, Department of Biology, Iowa City, IA 52242.

## DECLARATION OF INTEREST

The authors declare that they have no known competing financial interests or personal relationships that could have appeared to influence the work reported in this paper.

## AUTHOR CONTRIBUTIONS

**Xinyue Lu:** Conceptualization, Methodology, Validation, Formal analysis, Investigation, Writing – original draft, Writing – review & editing, Visualization. **Olga L. Sharko:** Methodology, Validation, Resources, Writing – review & editing. **Vadim V. Shmanai:** Methodology, Validation, Resources, Writing – review & editing. **Mikhail S. Shchepinov:** Conceptualization, Resources, Writing – review & editing, Supervision, Project administration. **James F. Markworth:** Conceptualization, Methodology, Software, Formal analysis, Resources, Writing – review & editing, Visualization, Supervision, Project administration, Funding acquisition.

## FUNDING

This work was supported by the U.S. Department of Agriculture National Institute of Food and Agriculture [Research Capacity Fund (HATCH Multistate), project no. 7004451 (NC1184)]; the National Institute of Diabetes and Digestive and Kidney Diseases of the National Institutes of Health [grant number R01DK132819]; and laboratory start-up funding provided by the Purdue University College of Agriculture to James F. Markworth. The funders had no role in study design, data collection and analysis, decision to publish, or preparation of the manuscript. The content is solely the responsibility of the authors and does not necessarily represent the official views of the National Institutes of Health, the USDA, or the U.S. Government. Any opinions, findings, conclusions, or recommendations expressed in this publication are those of the author(s) and should not be construed to represent any official USDA or U.S. Government determination or policy.

## Notes

### Competing Interest Statement

The authors have declared no competing interest.

